# The impact of microclimate and soil on the ecology and evolution of an arctic plant

**DOI:** 10.1101/2020.07.26.221721

**Authors:** Niklas J. Wickander, Pil U. Rasmussen, Bryndís Marteinsdóttir, Johan Ehrlén, Ayco J. M. Tack

## Abstract

The arctic and alpine regions are predicted to experience one of the highest rates of climate change, and the arctic vegetation is expected to be especially sensitive to such changes. Understanding the ecological and evolutionary responses of arctic plant species to changes in climate is therefore a key objective. Geothermal areas, where temperature gradients naturally occur over small spatial scales, and without many of the confounding environmental factors present in latitudinal and other gradient studies, provide a natural experimental setting to examine the impact of temperature on the response of arctic-alpine plants to increasing temperatures. To test the ecological and evolutionary response of the circumpolar alpine bistort (*Bistorta vivipara*) to temperature, we collected plant material and soil from areas with low, intermediate, and high soil temperatures and grew them in all combinations at three different temperatures. At higher experimental soil temperatures, sprouting was earlier, and plants had more leaves. Sprouting was earlier in soil originating from intermediate temperature and plants had more leaves when grown in soil originating from low temperatures. We did not find evidence of local adaptation or genetic variation in reaction norms among plants originating from areas with low, intermediate, and high soil temperature. Our findings suggest that the alpine bistort has a strong plastic response to warming, but that differences in soil temperature have not resulted in genetic differentiation. The lack of an observed evolutionary response may, for example, be due to the absence of temperature-mediated selection on *B. vivipara*, or high levels of gene flow balancing differences in selection. When placed within the context of other studies, we conclude that arctic-alpine plant species often show strong plastic responses to spring warming, while evidence of evolutionary responses varies among species.

## Introduction

Warming in the Arctic is predicted to be particularly strong (Pachauri et al. 2015). This increase in temperature may have a major impact on the phenology, growth and reproduction of arctic plants, whose life cycles are strongly influenced by seasonal shifts in temperature (Henry and Molau 1997, Callaghan et al. 2004, Gornish and Prather 2014, Semenchuk et al. 2015). To predict the long-term effects of increasing temperatures on the arctic flora, it is important to understand both immediate and evolutionary responses. For example, plant individuals may respond plastically and advance development in response to increasing spring temperatures (Anderson et al. 2012). At the same time, overall increases in temperatures might lead to changes in how strong the optimal response of development to temperature should be, and to evolutionary responses in reaction norms (Valdés et al. 2018). To study the response of arctic plants to increasing temperature, studies commonly use experiments with open-top climate chambers (Arft et al. 1999, Wolkovich et al. 2012), or sample plants growing along altitudinal (Frei et al. 2014) and latitudinal gradients (Toftegaard et al. 2016). Such methods have limitations, with the time-scale of experimental work often too short to observe evolutionary responses, and changes in temperature along altitudinal and latitudinal gradients often occurring alongside changes in other environmental factors. Given these limitations, geothermal systems, where radiation heating causes small scale variation in temperature but not in other environmental variables (O’Gorman et al. 2014, Sigurdsson et al. 2016), offer an alternative setting where plastic and evolutionary responses to temperature can be explored (Gudmundsdottir et al., 2011; O’Gorman et al., 2014; Sigurdsson et al., 2016; Valdés et al., 2018). For example, Iceland has many naturally occurring geothermal areas, where volcanic activity close to the surface warms water and soil through radiation heating (Gudmundsdottir et al. 2011, Zakharova and Spichak 2012). In such geothermal areas, plants of the same species frequently grow side-by-side in heated and non-heated soil.

Plasticity allows plants to grow under variable conditions and to respond to changes in the environment (Agrawal 2001). For example, plants often germinate earlier and grow larger at higher temperatures (Anderson et al. 2012). The optimal plastic response of plants is likely to differ depending on their local environmental conditions (Grether 2005). If there is genetic variation in plasticity, we would thus expect selection to result in reaction norms that are locally adapted (Valdés et al. 2018). For example, plants that have evolved in a low-temperature environment may rapidly germinate, flush leaves or flower in response to warming in spring, whereas plants from a high-temperature environment may respond more slowly to spring warming. Indeed, previous studies have found that plants from geothermally heated areas have an earlier spring phenology than plants from non-heated areas in the field, while the pattern is reversed when grown in a common environment, a phenomenon referred to as counter-gradient variation (Conover and Schultz 1995, Valdés et al. 2018).

The plastic responses and local adaptation of plants in geothermal landscapes may not only be in response to the direct effect of warming, but also due to indirect effects of warming on the local environment. One such potentially important indirect effect is the soil environment (Wittenmayer and Merbach 2005, Bardgett and van der Putten 2014). There is evidence for plant local adaptation to both abiotic (Bradshaw 1952, Joshi et al. 2001, Brady et al. 2005) and biotic soil characteristics (Mursinoff and Tack 2017). Warmer soils often have higher nutrient levels, which are frequently limited in the arctic environment (Sigurdsson et al. 2016). Moreover, both beneficial and antagonistic soil biota, like fungi, bacteria and archaea, might be more diverse in warmer soils (Tedersoo et al. 2014, Rasmussen et al. 2018, 2020).

The aim of this study was to investigate the effects of soil heating on plastic and evolutionary responses of the plant *Bistorta vivipara.* For this, we carried out a three-way multifactorial greenhouse experiment where we grew bulbils (i.e. asexually produced axillary buds), collected from locations with low, intermediate and high temperatures, in soils from the same three types of locations, under low, intermediate and high soil temperatures in a climate chamber. We tested the following specific hypotheses:

1. Experimental heating of soil results in a higher probability of sprouting, earlier sprouting, and increased size
2. The bulbils planted in soils originating from heated areas have a higher probability of sprouting, sprout earlier, and grow larger due to higher availability of mineralized nutrients
3. Selection has resulted in a counter-gradient pattern where bulbils originating from heated sites show a weaker growth response to temperature than bulbils originating from non-heated sites
4. Genetic differences among plants originating from different soil temperatures correspond to local adaptation, i.e. plants perform better under environmental conditions that correspond to conditions at their site of origin
5. Both the abiotic and biotic soil environment influence the ecological and evolutionary response of plants to heating

## Material and methods

### Study system

Geothermal landscapes, where magma gets close to the surface and heats soils and ground water, can act as hotspots of local adaptation (O’Gorman et al. 2014). Notably, within these landscapes the temperature can vary strongly at small spatial scales, with temperatures differing by several or tens of degrees Celsius between locations separated by only a few meters (Gudmundsdottir et al. 2011). The plant and soil microbial communities in these locations are frequently strongly affected by spatial variation in soil temperature (Friberg et al. 2009, Zakharova and Spichak 2012). At the same time, variation in soil nutrient concentrations across the temperature gradient is relatively limited, yet availability increases with rising temperatures in arctic ecosystems through mineralization of locked nutrients (Weintraub and Schimel 2003, O’Gorman et al. 2014, Semenchuk et al. 2015, Sigurdsson et al. 2016).

The alpine bistort (*Bistorta vivipara*) is a perennial herb that occurs in many parts of the northern hemisphere (Jonsell and Karlsson 2000). In Iceland, *B. vivipara* can be found growing naturally in a wide range of temperature conditions (Kristinsson 2001), including geothermal areas. The herb primarily spreads clonally by bulbils, and only rarely produces fertile seeds (Jonsell and Karlsson 2000). Bulbils are asexually produced bulb-like structures that replace the lower inflorescence of *B. vivipara*. Bulbils have no specialized dispersal structures, and often start growing within tens of centimeters from the mother plant (Bills et al. 2015).

### Origin of plants and soil

We collected plants and soil from Hengladalir, a geothermally active valley located a few tens of kilometers east of Reykjavik, Iceland. Within this area, we identified three locations with no heating (soil temperatures *c.* 13°C), three locations with slight heating (14-17°C) and three locations with strong heating (25-30°C). From each location, we collected soil up to a depth of *c.* 10 cm, which matches the average root depth of *B. vivipara*. From the soil, we removed the litter layer and larger rocks and roots. From each location, we further collected bulbils from one to three *B. vivipara* plants (henceforth referred to as ‘mother plants’), resulting in seven mother plants from each temperature.

### Experimental design

To investigate the effect of temperature and soil on plant performance, we designed a multifactorial experiment where bulbils collected at different temperatures (3 levels) were grown in soils collected at different temperatures (3 levels), and the soils were then experimentally heated to low, intermediate and high temperatures, to simulate the natural variation in soil temperature at the geothermal area. This resulted in 27 different treatment combinations. For each treatment combination, we had seven 82 mm diameter pots (330 ml) planted with two bulbils. We distributed bulbils from each mother plant equally across treatment combinations. As maternal investment in bulbils may affect growth, we recorded bulbil weight prior to planting.

We experimentally heated the plant pots in water baths. The water baths were uniformly heated using a combination of aquarium heaters (Easyheater 75W, AQUAEL, Poland) and water circulation pumps (Compact 300, EHEIM, Germany). In total there were nine water baths, three at each temperature level: i) low temperature (13°C), ii) intermediate (18°C) and iii) high (30°C). In each water bath, we placed 21 plant pots. Water baths were placed in a climate chamber set to 13°C and a day length of 16 h.

To avoid microbial contamination, we sealed the plant pots, but filled the lower 2.5 cm with gravel and water-absorbing cloth to allow for drainage. To allow water to flow below the plant pot, we glued three metal washers under each plant pot. To avoid positional effects, the location of the plant pots was regularly randomized within the three water baths belonging to the same temperature treatment.

To disentangle the effect of soil biota from that of the abiotic soil environment (e.g. physical or chemical structure) on the probability and timing of bulbil sprouting, we set up a complementary experiment using sterile soil. For this, we applied only the low and high temperatures, resulting in eight treatment combinations and 56 plant pots. For sterilization, soil was autoclaved for 1 h at 121°C.

### Plant performance measures

Plants were monitored every other day for sprouting. The day of sprouting was determined as the number of days from planting until the first sign of sprouting was observed. Plant height and number of leaves were recorded on days 84 and 126, and the length and width of the longest leaf were measured on days 97 and 159. Given the elliptical shape of the leaf, we calculated leaf size using the formula for the area of an ellipse:

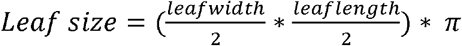

### Analysis

Data were analyzed using the generalized linear mixed model framework as implemented in R v. 3.6.1 (R Core Team 2019). We tested for significance using the *Anova* function in the *car* package (Fox and Weisberg 2019).

To test for the impact of soil temperature and origin of plants and soil on bulbil sprouting and plant growth, we modelled sprouting (0/1), day of sprouting, plant height, number of leaves and leaf size as a function of original soil temperature, original bulbil temperature, experimental soil temperature and their two- and three-way interactions. Bulbil weight was included as a covariate. To account for variation among mother plants, we included the random factor ‘*Mother plant*’. As we planted two bulbils in each plant pot, we further included the random factor ‘*Plant pot*’ which was nested under ‘*Mother plant*’. For probability of sprouting, we used a binomial distribution with a logit link; for day of sprouting, plant height and leaf size, we used a normal distribution with an identity link; and for the number of leaves, we used a Poisson distribution with a log link.

We *a priori* planned to test for local adaptation in those cases where we found an interaction between original bulbil temperature and original soil temperature, or between original bulbil temperature and experimental soil temperature (Kawecki and Ebert 2004, Blanquart et al. 2013), using a contrast statement as implemented with the function *emmeans* in the R-package *emmeans* (Lenth 2019).

An effect of temperature of the soil at the collection site on the probability of sprouting and the date of sprouting could be due to either the abiotic or biotic soil characteristics. To disentangle the role of the soil abiotic and biotic environment, we first created a data set with data from both field and sterile soil experiment, where we only retained those treatment combinations present in both experiments (i.e. only ‘low’ and ‘high’ temperatures). We then modelled the affected plant traits as a function of original soil temperature, original bulbil temperature, experimental soil temperature and soil sterilization, including all two- and three-way interactions. As above, we included the random factors ‘*Mother plant*’ and ‘*Plant pot*’ (nested under ‘*Mother plant*’). Given the many terms in the full model, we then identified the minimum adequate model using backward selection, using a p-value of 0.05 as a threshold (Crawley 2013).

## Results

Bulbils were more likely to sprout and sprouted earlier when grown at higher soil temperatures (Fig 1AB, Table 1). Plants had the smallest leaves when grown at intermediate soil temperature when measured 97 days after planting, whereas plants grown at higher soil temperatures had more leaves when plants were measured 126 days after planting (Fig. 1CD, Table 1). Bulbils sprouted earlier in soil originating from areas with intermediate temperature than in soil originating from low or high temperatures (Table 1, Fig. 2A). Plants had a higher number of leaves at 126 days after planting when grown in soil from a low temperature origin (Fig. 2B, Table 1). At day 159, the end of the experiment, leaf size was interactively affected by experimental soil temperature and original soil temperature: plants had larger leaves when grown in soil originating from intermediate temperature, but only when the experimental soil temperature was either low or high (Fig. S1). Plant height increased with bulbil weight, but was not affected by any of the experimental factors (Table 1).

**Table 1.** The impact of original soil temperature (OST), original bulbil temperature (OBT) and experimental soil temperature (EST), as well as lbil weight, on the performance of *Bistorta vivipara* in field soil. Shown are P-values, with significant values in bold. The number of replicates each response variable is given within parentheses.

|  | Original soil temperature<br>(OST) | Original bulbil<br>temperature (OBT) | Experimental soil<br>temperature (EST) | Bulbil weight | OST × OBT | OST × EST | OBT × EST | OST × OBT × EST |
| --- | --- | --- | --- | --- | --- | --- | --- | --- |
| Sprouting success (n = 378) | 0.856 | 0.849 | **<0.001** | 0.164 | 0.121 | 0.212 | 0.276 | 0.313 |
| Day of sprouting (n = 240) | **<0.001** | 0.094 | **<0.001** | 0.072 | 0.935 | 0.080 | 0.456 | 0.445 |
| Plant height on day 84 (n = 217) | 0.792 | 0.088 | 0.338 | **<0.001** | 0.790 | 0.106 | 0.753 | 0.468 |
| Plant height on day 126 (n = 77) | 0.401 | 0.648 | 0.529 | **0.030** | 0.561 | 0.337 | 0.623 | 0.102 |
| Leaf number on day 84 (n = 217) | 0.291 | 0.514 | 0.177 | 0.561 | 0.874 | 0.785 | 0.816 | 0.992 |
| Leaf number on day 126 (n = 77) | **0.012** | 0.118 | **0.004** | 0.984 | 0.939 | 0.772 | 0.559 | 0.999 |
| Leaf size on day 97 (n = 162) | 0.066 | 0.296 | **0.043** | 0.067 | 0.538 | 0.652 | 0.050 | 0.571 |
| Leaf size on day 159 (n = 48) | 0.162 | 0.924 | 0.055 | 0.521 | 0.296 | **0.025** | 0.074 | - |

**Figure 1.**
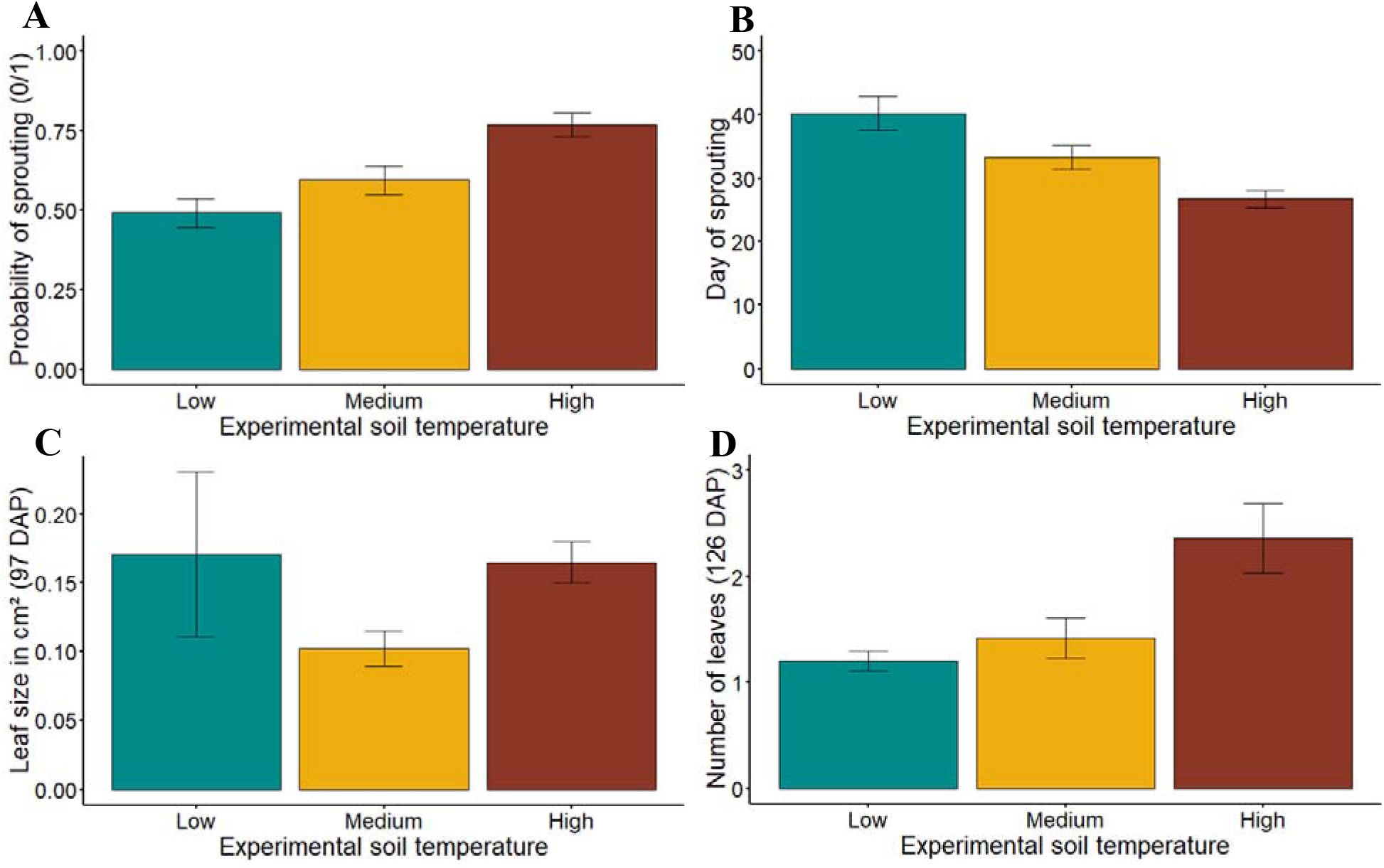
The impact of experimental soil temperature on bulbil sprouting and performance of *Bistorta vivipara*. Shown are the impact of experimental soil temperature on A) probability of sprouting, B) day of sprouting, C) leaf size at 97 days after planting (DAP) and D) number of leaves at 126 days after planting. Shown are means ± standard errors.

**Figure 2.**
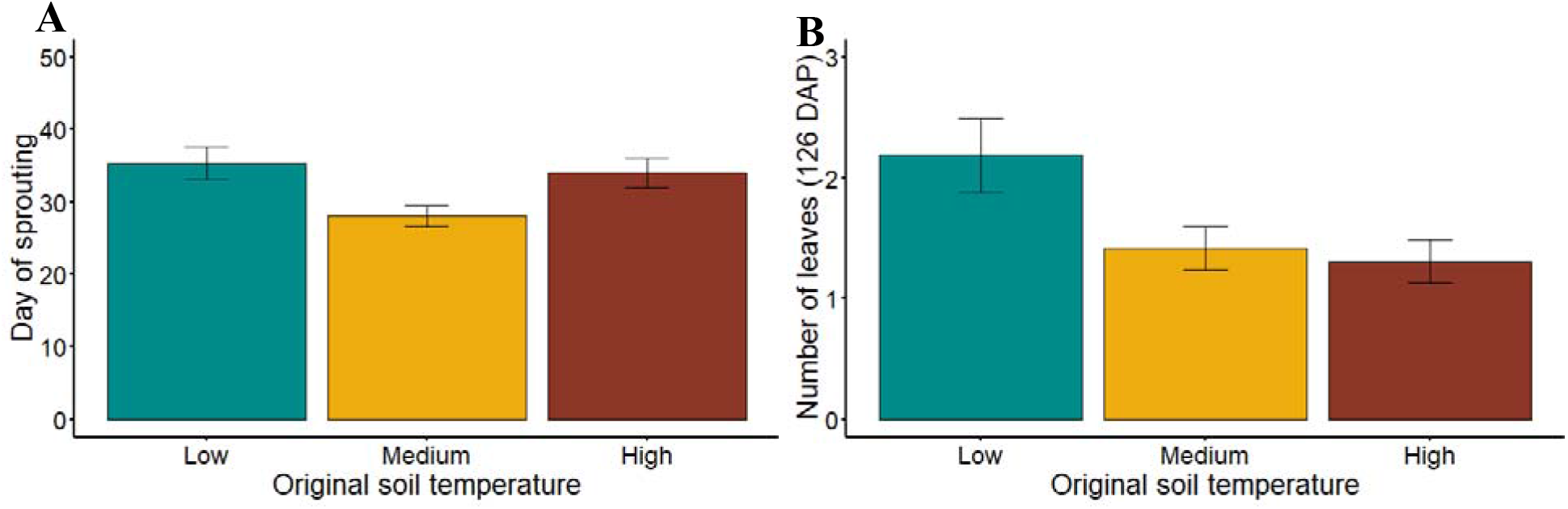
The impact of original soil temperature on bulbil sprouting and performance of *Bistorta vivipara*. Shown are the impact of experimental soil temperature on A) day of sprouting and B) the number of leaves at 126 days after planting (DAP). Shown are means ± standard errors.

Plants originating from areas with low, intermediate, and high soil temperature did not differ in probability of sprouting, day of sprouting or growth traits (Table 1). We also did not detect any evidence of local adaptation in terms of an effect of the interaction between original bulbil temperature and original or experimental soil temperature on sprouting probability, day of sprouting or plant growth (Table 1).

Bulbil sprouting was later in the sterile soil than in the field soil, and sprouting was earlier with experimental soil heating (Fig. S2, Table S1). We further detected a weak interactive effect between original bulbil temperature, experimental soil temperature and soil sterilization, suggesting that bulbils of a high temperature origin sprouted later than expected when growing in soil originating from an area with low temperature, but only when the soil was sterilized.

## Discussion

We used a multi-factorial experiment to investigate the impact of microclimate and soil on the immediate and evolutionary response of the alpine bistort. As expected, higher experimental soil temperature advanced bulbil sprouting and led to plants with more leaves. In contrast to our prediction, sprouting was earliest in soil originating from locations with intermediate temperature, and the number of leaves was highest when plants were grown in soil from a low temperature origin. We found no evidence for either genetic variation in phenotypic plasticity or local adaptation. The microbial community in the soil advanced bulbil sprouting, with some weak evidence for delayed sprouting of plants originating from high temperature soils when they were grown at low soil temperature in the absence of soil biota.

Plants sprouted earlier and had more leaves with higher experimental soil temperature. This is consistent with previous studies that have shown an advance of spring phenology and flowering, as well as increased plant growth, with increasing temperature (Arft et al. 1999, Valdés et al. 2018). For example, flowering phenology has been shown to advance with heating in geothermal areas (Anderson et al. 2012, Valdés et al. 2018), and plant biomass is higher in *Arctagrostis latifolia* and *Carex bigelowii* when grown at higher soil temperature (Marchand et al. 2005). Taken together, these findings suggest that the circumpolar *B. vivipara* will start growing earlier, and grow larger, with climate change.

Surprisingly, the bulbils sprouted earliest in soil from the intermediate temperature origins, and plants had the highest number of leaves when grown in soil from low temperature origins. This did not match our expectation that the higher mineralization rates, and associated higher availability of nutrients, in soils from high temperature would result in earlier sprouting and increased performance. Previous studies have shown that nutrients are frequently a limiting factor in arctic ecosystems and increased temperatures enhance nutrient availability for plants and increase growth (Henry and Molau 1997, Weintraub and Schimel 2003, Semenchuk et al. 2015). Moreover, warmer soil is often associated with a more diverse soil community and higher soil biotic activity (Friberg et al. 2009, Zakharova and Spichak 2012). One explanation for the lack of the expected pattern is that nutrient variation across temperature gradients is limited in geothermal areas (O’Gorman et al. 2014, Sigurdsson et al. 2016), and even if mineralization is higher in soils in heated areas, this may not result in higher availability of nutrients when the soil is exposed to lower temperatures. Importantly, our findings illustrate that warming-mediated changes in the soil environment can have idiosyncratic effects on plant phenology and plant traits.

We found no evidence for genetic variation in phenotypic plasticity or local adaptation of plants to spatial variation in soil temperature. This contrasts to previous studies, which found adaptive patterns of phenotypic plasticity by comparing plants grown in the field and common garden (Valdés et al. 2018), and local adaptation using reciprocal transplant experiments (Conover and Schultz 1995, Joshi et al. 2001, Anderson et al. 2012). For example, Valdés et al. (2018) found that the arctic plant *Cerastium fontanum* flowered earlier when growing in heated areas than in non-heated areas in the field, but that individuals from heated areas flowered later than individuals from non-heated areas when growing in a common garden. The lack of counter-gradient variation and local adaptation in *B. vivipara* might have three non-mutually exclusive explanations. First, selection on the investigated traits may be weak or absent. Second, *B. vivipara* mostly reproduces clonally, and the lack of recombination may slow down the rate of local adaptation compared to sexually reproducing plant species. Indeed, sexually reproducing species used in other studies showed adaptive patterns of counter-gradient variation (Anderson et al. 2012, Valdés et al. 2018). Finally, gene flow may swamp the effect of natural selection, but as dispersal distances are low in *B. vivipara*, this appears less likely to explain the absence of an evolutionary response to temperature in this species. To tease apart these explanations, future studies could use quantitative genetic approaches to assess additive genetic variation for the investigated traits, as combined with a genetic or experimental assessment of gene flow.

Interestingly, bulbil sprouting was slower in the sterile soil than in the field soil, indicating that the soil microbial community stimulates bulbil sprouting. The effect of soil biota matches previous studies, which found that soil organisms can advance phenology and influence plant growth (Wagner et al. 2014, Rasmussen et al. 2017, 2020). The delayed sprouting of bulbils originating from high temperature when grown at low soil temperature, but only when the soil was sterilized, may indicate that soil biota plays a role in the evolutionary response of plants to soil heating. However, we have no insights into the putative adaptive nature of this pattern.

## Conclusions

Our findings highlight that the circumpolar alpine bistort has a strong plastic response to temperature. We did not find an evolutionary response in the form of counter-gradient variation or local adaptation. This contrasts with previous reports from other plant species in geothermal areas, and along latitudinal and altitudinal gradients, and may imply that the relative importance of plastic vs genetic responses is highly variable among plant species. Such variable responses, in combination with underlying mechanisms, have profound implications for our understanding of how the arctic vegetation will respond to a changing climate. For example, some plant species may lack genetic variation for selection to act on, and thereby show maladaptive plastic responses to changes in temperature. Other plant species may have additive genetic variation, but not show an evolutionary response in the current context due to high rates of gene flow. Still other plant species may have a relatively weak plastic response but show local adaptation across small spatial scales. Overall, we hope that future studies will test for the ecological and evolutionary response of plants to changes in temperature across a broad range of plant species and environmental clines, and unravel the underlying mechanisms, which will allow for the development of eco-evolutionary models to predict the response of the arctic vegetation to changes in climate. This will give insights in the threats posed by climate change to the arctic vegetation and help inform about potential mitigation actions.

## Declarations

We thank Jordbruksverket for the import permit (6.5.18-5502/16) for the soil and plant material used in this study. We thank Nia Sigrún Perron for help in the field. This research was supported by a grant from the Swedish Research Council (2015-03993 to AJMT) and funding from the University of Iceland post◻doctoral grant to B.M. The authors declare no conflict of interest. NJW, PUR, JE and AJMT designed the experiment. BM collected field materials, and NJW carried out the experiment and analyzed the data. NJW wrote the manuscript, with contributions from all authors.

## Notes

### Competing Interest Statement

The authors have declared no competing interest.

