## Supplemental Tables and Figures for "The impact of microclimate and soil on the ecology and evolution of an arctic plant"

**Supplementary material**

**Table S1.** The impact of original soil temperature (OST), original bulbil temperature (OBT), experimental soil temperature (EST) and soil sterilization (FS), as well as bulbil weight, on the timing of sprouting of bulbils of *Bistorta vivipara*. Shown are P-values from the minimum adequate models (where original soil temperature was excluded), with significant values in bold.

|  | Day of sprouting |
| --- | --- |
| Original bulbil temperature (OBT) | 0.524 |
| Experimental soil temperature (EST) | **<0.001** |
| Field or sterile (FS) | **<0.001** |
| Bulbil weight | 0.293 |
| OBT × EST | 0.389 |
| OBT × FS | 0.418 |
| EST × FS | 0.127 |
| OBT × EST × FS | **0.049** |

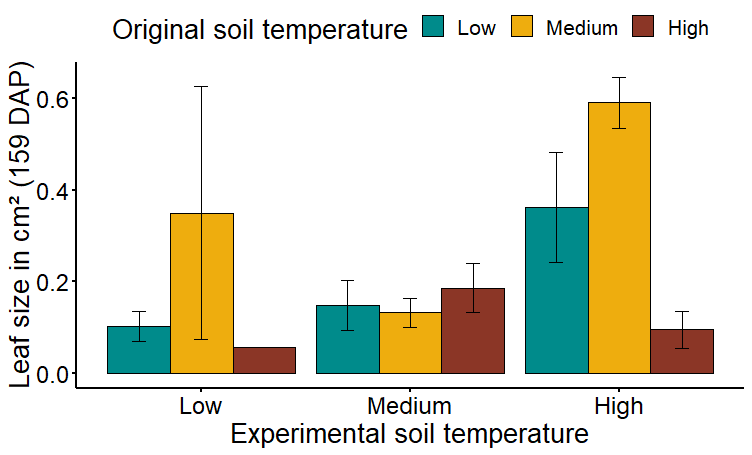
**Figure S1** The influence of original soil temperature and experimental soil temperature on leaf size of *Bistorta vivipara* 159 days after planting (DAP). Shown are means ± standard errors.

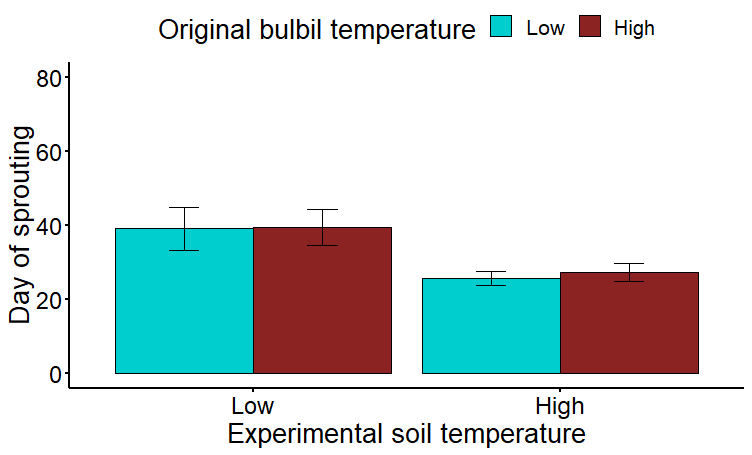

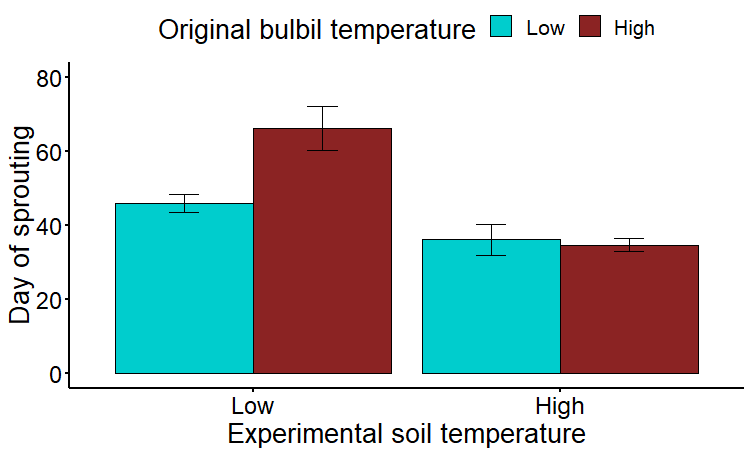

**A**

**B**

**Figure S2** The impact of original bulbil temperature and experimental soil temperature on the day of sprouting of bulbils of *Bistorta vivipara*, separately for plants grown in A) field and B) sterile soil. Shown are means ± standard errors.
